# Learning and Mapping Lyme Disease Patient Trajectories from Electronic Medical Data for Stratification of Disease Risk and Therapeutic Response

**DOI:** 10.1101/239020

**Authors:** Osamu Ichikawa, Benjamin S. Glicksberg, Brian Kidd, Li Li, Joel T. Dudley

## Abstract

**Background:** Lyme disease (LD) is an epidemic, tick-borne illness with approximately 329,000 incidences diagnosed each year in United States. Long-term use of antibiotics is associated with serious complications, including post-treatment Lyme disease syndrome (PTLDS). The landscape of comorbidities and health trajectories associated with LD and associated treatments is not fully understood. Consequently, there is an urgent need to improve clinical management of LD based on a more precise understanding of disease and patient stratification.

**Methods:** We used a precision medicine machine-learning approach based on high-dimensional electronic medical records (EMRs) to characterize the heterogeneous comorbidities in a LD population and develop systematic predictive models for identifying medications that influence the risk of subsequent comorbidities.

**Findings:** We identified 3, 16, and 17 comorbidities at broad disease categories associated with LD within 2, 5, and 10 years of diagnosis, respectively. At higher resolution of ICD-9 levels, we pinpointed specific co-morbid diseases on a timescale that matched the symptoms associated with PTLDS. We identified 7, 30, and 35 medications that influenced the risks of the reported comorbidities within 2, 5, and 10 years, respectively. These medications included six previously associated with the identified comorbidities and 29 new findings. For instance, the first-line antibiotic doxycycline exhibited a consistently protective effect for typical symptoms of LD, including ‘backache Not Otherwise Specified (NOS)’ and ‘chronic rhinitis’, but consistently increased the risk of ‘cataract NOS’, ‘tear film insufficiency NOS’, and ‘nocturia’.

**Interpretation:** Our approach and findings suggest new hypotheses for precision medicine treatments regimens and drug repurposing opportunities tailored to the phenotypic profiles of LD patients.

**Funding:** The Steven & Alexandra Cohen Foundation

## INTRODUCTION

Lyme disease (LD) is a vector-borne, infectious disease caused by the bacterium *Borrelia burgdorferi* that is transmitted to humans through tick bites. According to the US Centers for Disease Control and Prevention (CDC), around 329,000 LD cases occur annually and it becomes a major US public health problem that causes substantial use of health care resources. LD is most prevalent in the Northeast and upper Midwest, and 95% of all confirmed cases in 2015 were reported in 14 states ^1^. The symptomology of LD is heterogeneous, although some general patterns have emerged. The first manifestation of LD is often an expanding annular lesion, called erythema migrans, near the bite location, but this sign is present in only 70–80% of patients ^2^. The length of time for the rash to occur, along with the characteristics of the rash (e.g., composition and size) can also vary ^3^. Other clinical features that often arise, singly or in combination, include fever, pain, arthritis, myopericarditis, neurological symptoms (e.g., facial nerve palsy), and satellite rashes. One explanation for this variability is that the genotype of the tick itself might affect aspects of pathogenesis, such as the probability of hematogenous dissemination ^4,5^. The neurological manifestations in LD, reported in 3–12% of patients, are of greatest concern ^6^. These phenomena, collectively called neuroborreliosis, are often associated with intense pain that can manifest either soon after infection or much later, from months to years afterward.

Accurate and precise diagnoses of LD present several challenges. Typically, laboratory testing of LD follows identification of cutaneous manifestations from visual inspection but these manifestations are not always present. Current guidelines recommend serologic testing, a two-phase process consisting of an enzyme immunoassay within 30 days of symptom onset, followed by Western blot after 30 days from symptom occurs ^7,8^ if the early test is active. Even together, this diagnostic strategy has poor sensitivity, particularly during the acute phase, with false-negative rates of up to 50% ^9^. Other laboratory methods are specific for particular manifestations, e.g., testing of CSF for central nervous system involvement. Recent work has shown that incorporation of data from various wearable devices can detect early signs of LD and associated inflammatory responses. For example, variations in peripheral capillary oxygen saturation (SpO2), a marker associated with physiological macro-phenotypes such as fatigue, can be measured by portable biosensors to facilitate more accurate and rapid LD diagnosis from variations in these measurements and could be economically feasible for widespread use in the future. Currently, however, clinicians still have to rely on traditional measures to diagnose patients^10^. Furthermore, comorbid conditions can interfere with both diagnosis and treatment. For instance, other infections can be concurrently transmitted with LD ^11^, making differential diagnosis even more difficult and sometimes requiring specialized, alternative treatment strategies. Many studies have attempted to develop methods for differentiating LD from other similar syndromes, e.g., septic arthritis vs. LD of the knee in children ^12^.

Following successful diagnosis, LD is most commonly treated with antibiotics such as doxycycline, amoxicillin, cefuroxime axetil, and ceftriaxone. Although these medications have high cure rates (~90%) ^13^, they are associated with serious complications and adverse events, especially under prolonged use ^14,15 3,16^. Notably in this regard, one study showed that certain first-line treatments, specifically intravenous ceftriaxone followed by doxycycline for chronic LD patients were not effective compared to placebo forcing discontinuation of the trial ^17^. Another study reported that repeated IV ceftriaxone treatment for Lyme encephalopathy resulted in only minor cognitive improvements, with high rates of relapse of cognitive symptoms ^18^. These findings suggest that unknown factors are responsible for the high variability of treatment outcomes for patients with LD. Additionally, up to 20% of treated patients develop posttreatment Lyme disease syndrome (PTLDS), in which lingering symptoms such as fatigue, pain, or joint and muscle aches last for months or even years. The causes and frequencies of these symptoms remain unclear, and the issue is further confounded by the presence of concurrent diseases.

Several recent studies reveal the uncharacterized complexity of disease course and treatment response in LD. One study found certain first-line treatments, specifically intravenous ceftriaxone followed by doxyclycline, for chronic LD patients were not effective compared to placebo and were forced to discontinue their study^17^. Another study assessed the effects of repeated IV ceftriaxone treatment for Lyme encephalopathy and found only minor cognitive improvements with high rates of relapse in cognition issues ^18^. Therefore, it seems that there are other factors at play that may explain the high variability of treatment outcomes for patients with LD. It is difficult to disentangle to what extent given treatment responses and disease sequelae are due to differences in individual immune responses, patient characteristics, disease burden, and treatment timing, or to the medications themselves. Indeed, it is very likely that response and outcome depend on a complex interplay between these factors, making clinicians’ jobs extremely difficult. To address the diverse symptomology, imperfect diagnostic strategies, and variable treatment outcomes of LD, comprehensive study designs are required. For example, an investigation of risk factors for LD infection, such as behavioral and environmental risk factors, revealed that LD–positive serology is significantly associated with clinical and demographic features such as previous self-reported LD diagnosis and age, behavioral factors such as wearing protective clothing, and geographic/environmental factors such as shrub edge density in property location ^19^. Another study evaluated the risks to individuals based on geographical features such as the suitability of the local habitat for ticks ^20^.

Although the aforementioned studies have provided a great deal of useful information, the variability in global risk profiles for LD pathogenesis remains incompletely understood, and there is an unmet need for personalized treatment recommendations that take into account individual characteristics such as demographics and disease burden. Electronic Medical Records (EMRs) from hospitals contain a wealth of longitudinal, patient-level data encompassing prior history of prescriptions and disease diagnoses, along with clinical outcomes, that can be exploited to investigate these issues in a data-driven fashion. To date there has been no systematic analysis of LD using EMR data, particularly from a hospital within a high-risk state. In this study, we leverage an EMR data set representing over five million unique patients of diverse racial and ethnic backgrounds collected from a large academic medical center in New York City. Although not itself located adjacent to a wooded area, MSH caters to patients from all over the state. New York is one of the aforementioned 14 states reporting the vast majority of LD cases, and in 2017 had an incidence rate of 16.4 per 100,000 individuals (CDC), one of the highest in the country.

We hypothesized that EMR data from MSH could provide a rich framework for studying the heterogeneity of Lyme manifestation, as well as the quality and efficacy of treatment. Using various state-of-the-art statistical and machine learning methods, we sought to identify patterns of clinical outcomes in order to help physicians develop treatment strategies tailored to patients’ disease profiles. Specifically, we investigated how demographic and clinical factors affect LD manifestations and clinical outcomes in the context of various treatments. Our main goals were to identify comorbidities associated with LD and develop a systematic predictive model for identifying medications that influence the risk of these conditions. As an alternative to one-size-fits-all strategies for treating LD, our methodology facilitates directing treatment recommendations and identifying possible repurposing opportunities tailored to the phenotypic profiles of Lyme patients. Our study is the first data-driven effort to prioritize medications for LD based on an individual’s phenotype profile. We identified Lyme-associated comorbidities at the level of broad disease categories, pinpointed specific co-morbid diseases associated with LD over time, and used machine learning to predict medications that influence the risks of these comorbidities.

We expect that the novel framework and findings from this study can support current and future efforts to develop personalized treatment strategies for patients with LD, including efforts to provide physicians with a broader evidentiary foundation on which to base their treatment recommendations (e.g., selection of antibiotics) based on individual patients’ disease risk profiles. Additionally, more precise knowledge of predicted adverse events would facilitate improvements in monitoring and management strategies.

## METHODS

### Patient population and standardization of clinical terminology

#### Patient cohort

We utilized Electronic Medical Records (EMRs) from the Mount Sinai Data Warehouse (MSDW), the largest comprehensive EMR system in New York City, which includes data from a racially and ethnically diverse patient base. Since 2000, more than 4.5 million unique patient records have been documented in this system. Disease diagnoses are encoded as International Classification of Diseases, 9th Revision (ICD-9) billing codes, which have been used extensively in EMR-related analyses ^21^. In this study, we retrieved records from all patients diagnosed with Lyme disease (LD) with the ICD-9 code 088.81 (*n*=2,134). We restricted the data to records occurring between 2000 and 2015, allowing for up to 15-year follow-up. In accordance with Health Insurance Portability and Accountability (HIPAA) and Protected Health Information (PHI) guidelines, the ages of these patients were censored at 18 and 90. Finally, we only kept data from patients with defined age, self-reported sex, and self-reported race/ethnicity (referred to as “race” in this manuscript). For the total of 1,767 Lyme patients, there were 930 females (52.6%) and 837 males (47.4%), with an average age of 47.8 ± 19.7. The racial breakdown of the cohort is as follows: 1,201 Caucasian (70.0%), 49 African-American (2.8%), 34 Hispanic/Latino (1.9%), and 483 Others (27.3%). For these patients, we also retrieved all other available clinical variables from EMR, including prescriptions and other disease diagnoses. In addition, we retrieved IgM/IgG lab measurements pertinent to LD. IgM or IgG Western blot labs were available for 28 patients after or at the time of diagnosis (45 days window of diagnosis). Of those, 89% patients (25/28) were reported as either IgM or IgG Western blot–positive, confirming true positivity for Lyme. The remaining 11% of patients (3/28) were reported as IgM-negative, but all were positive for p23, an antigen specific to Lyme. In total, we compiled 3,936 diseases diagnosis and 5,723 prescriptions. We provide a schematic of our study design, approach, and patient selection criteria in Figure 1.

**Figure 1.**
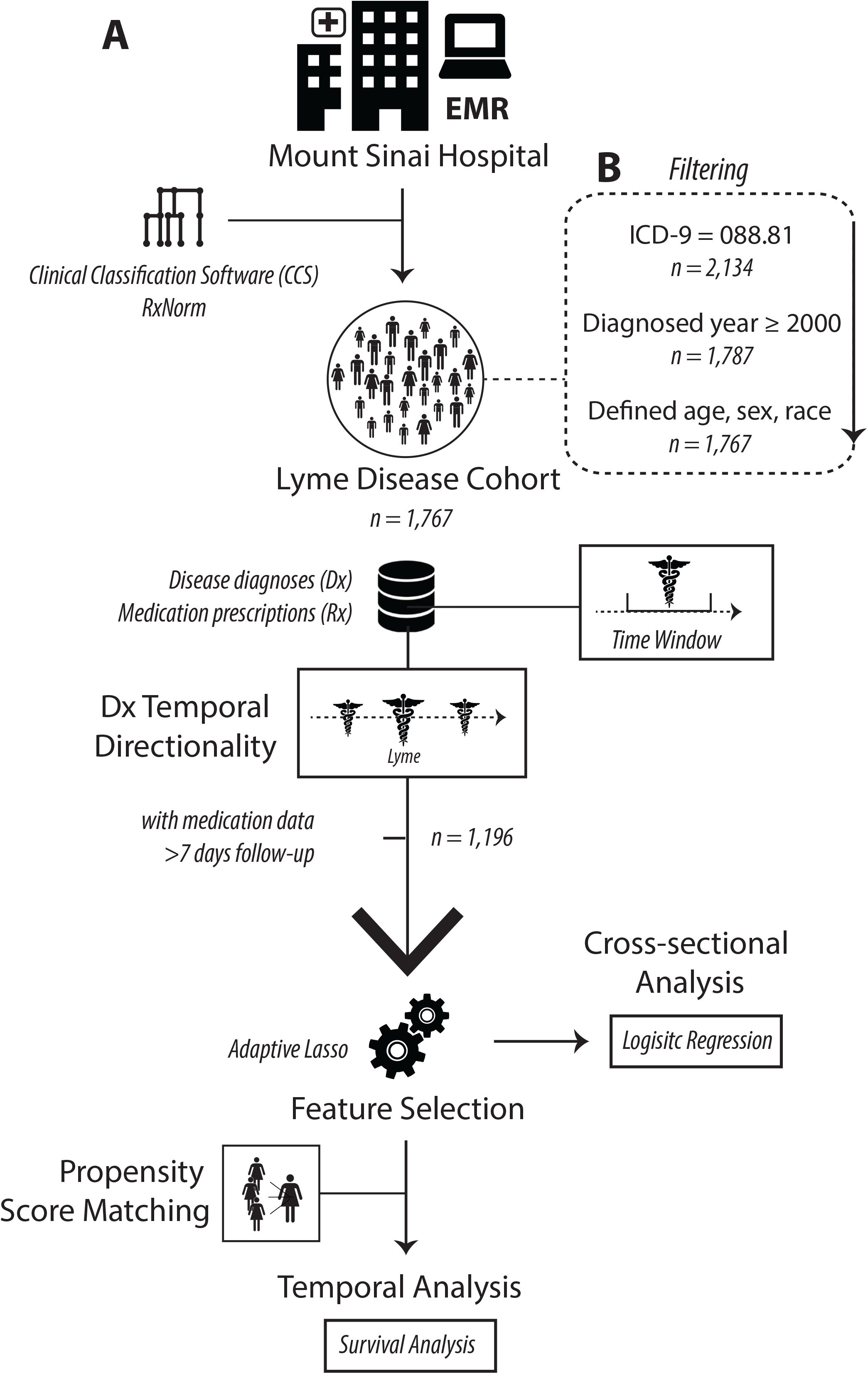
Workflow of the study, outlining steps from data organization to statistical methodologies.

#### Clinical sources and term standardization

We categorized diseases using the Clinical Classifications Software (CCS) for ICD-9 diagnosis codes, developed by AHRQ ^22^, which aggregates and characterizes more than 14,000 ICD-9 codes into broader coherent disease categories. This strategy helps to avoid sample size limitation as a result of using ICD-9 codes alone. For categorization, we used the ‘Single-Level Diagnosis’ (CCS-single) level, which has a total of 283 different categories. We standardized medication data by mapping to the RxNorm ontology ^23^. Specifically, we mapped these terms to ingredient codes, yielding 793 normalized medications. To ensure the robustness of our analyses, we required a sample size of > 20 patients for calculations of the significance of disease directionality and disease–medication association.

### Machine learning methods and analysis

#### Disease pair temporal directionality

For all patients with LD, we first assessed disease-pair connectivity patterns for comorbid diseases. Specifically, we determined whether the members of each pair exhibited a significant pattern in their temporal order, e.g., whether one preceded the other more often than expected by chance. We performed a cumulative binomial probability test to assess the temporal ordering of the associations between Lyme and all other diseases, assuming a 50% probability of either to occur before the other. We performed the following analysis on both broad and narrow disease categories. At the broader level, we analyzed representative CCS-single-level categories because this strategy could enhance signals that might be lost due to small sample size at the ICD-9 level. Second, we performed the analysis using standard ICD-9 codes in order to detect associations at a higher resolution for certain codes that may be more prevalent. Because these comorbid conditions can be either chronic or acute, we performed several iterations of this analysis over different time windows, specifically 2, 5, and 10 years.

For each time window, we restricted collection of information for the comorbid diseases in both temporal directions, relative to the date of first Lyme diagnosis. For the 2-year window, for example, we only collected disease data for each patient 2 years before and 2 years after the date of Lyme diagnosis. For the CCS-single- and ICD-9–level analyses, we performed 275 and 3,639 tests for each window, respectively. Last, to determine whether disease pairs with significant temporal directionality were also significantly comorbid, we performed a logistic regression for each pair controlling for age, sex, and self-reported race. The outcome variable in this model was the disease that was shown to occur after the other in the temporal analysis (significant in the binomial assessment).

#### Definition of outcomes and covariates in the machine-learning model

To discover risk factors or new therapeutic options for LD sequelae, we focused on the new onset of disease comorbidities more than 7 days after the diagnosis of LD. Of the 1,767 LD patients in the overall cohort, we systematically assessed the comorbidities and medication associations for 1,196 patients who were followed up for more than 7 days and had at least one prescription record in MSH’s EMR system. Like our disease-pair temporal directionality analysis, we set time windows of 2, 5, and 10 years. For each patient, we collected the diseases diagnosed within 2, 5, or 10 years after their first Lyme diagnosis date. We also retrieved medications prescribed within 1 year prior to and 2, 5, or 10 years after the first Lyme diagnosis. Outcome comorbidities were defined by ICD-9 code and categorized using CCS-single-level Diagnosis terms.

#### Feature selection

We considered many disease variables, coded by CCS-single-level categories, and medication variables, which were mapped to RxNorm ingredient codes. Accordingly, we adopted a feature selection method, penalized logistic regression with the adaptive LASSO (Eq. 1), to identify variables of the highest relevance that associated with ensuing comorbidities following LD diagnosis. The adaptive LASSO is an extension of the traditional LASSO ^24^ that uses coefficient-specific weights ^25^. The adaptive LASSO estimator selects the zero coefficients of the true parameters are estimated as zero with probability tending to one, which is called sparsity property. And the non-zero components are estimated as if the true sparse model were known *a* prior, which is asymptotically normal ^26^. Let 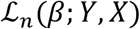 be the negative log-likelihood parametrized by β for a sample of size n. The adaptive LASSO estimator is defined as:

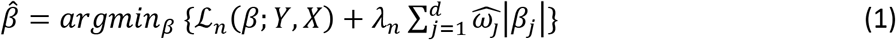

where 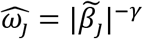 is a coefficient specific weights vector, and *λ_n_* is a regularization parameter. We set the positive constant *γ* as 1 according to Zou et al. ^25^, and obtained 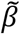 by the maximum likelihood estimate of Ridge regression. The *λ_n_* value for minimum AUC was chosen by 10-fold cross validation. We used the R package *glmnet* ^27^ for these penalized regressions.

#### Logistic regression model

We used odds ratio (OR) from logistic regression (Eq. 2) to assess the risk of future comorbidity progression on each medication taken (i.e. either increased risk or protective effect). We analyzed the pairs of outcome disease comorbidity and the medications that were selected by the adaptive LASSO. In this model, we adjusted for age, sex, self-reported race, and the follow-up time frame.

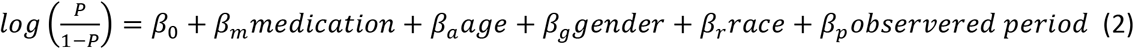

where *P* is the probability of a disease, medication is a binary variable, age is a continuous parameter, gender is a binary variable (Female/Male); race is a categorical variable (Caucasian, African American, Hispanic/Latino, or Other), and observed period is a continuous parameter. *β* coefficients for each covariate represent the effect size when controlling for all others.

#### Propensity score matching

To control for potential confounding factors due to imbalances of clinical characteristics, not limited to age and gender, we analyzed the temporal effects of medications after the propensity score matching to select an appropriate control cohort for the targeted case cohort ^28^. Thus, we created comparable cohorts, consisting of groups treated or untreated with a targeted medication, based on a set of covariates at the baseline time point, i.e., time zero for each patient. The baseline time point was defined as the first prescription day of the targeted medication or 7 days after LD diagnosis, whichever was later, because we observed disease comorbidities for more than 7 days after LD diagnosis.

The propensity scores of targeted prescriptions were predicted by a logistic regression model, including other significant medications and disease confounders selected by the adaptive LASSO with a 10-year time window, with patient demographics as covariates. Each patient prescribed a given medication was matched to a corresponding comparison patient (1:1 ratio) by nearest-neighbor matching. For instance, we analyzed association between doxycycline and ‘backache Not Otherwise Specified (NOS)’ (ICD-9 code: 724.5), ‘chronic rhinitis’ (472.0), ‘tear film insufficiency (insuffic) NOS (375.15)’, and ‘cataract NOS (366.9)’, and between amoxicillin and ‘acute upper respiratory infection (URI) NOS (465.9)’. A total of 328, 330, 358, 370, and 115 subjects were selected for each medication-comorbidity pair in the propensity score-matched treated/untreated group. The R package *MatchIt* ^29^ was used for propensity score matching.

#### Survival analysis

We generated survival curves by the Kaplan–Meier method and examined differences in survival among subgroups by the log-rank test, with propensity score matching of cases and controls. We calculated hazard ratios using Cox proportional hazards models:

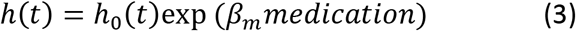

where *h*(*t*) is the expected hazard at time t, *h*_0_(*t*) is the baseline hazard, and medication is a binary variable. We verified the proportional hazards assumption by confirming that Schoenfeld residuals are independent of time (Schoenfeld test p > 0.1). We used the R packages *survival* and *survminer* for the survival analysis.

## RESULTS

### Identified comorbidities associated with Lyme disease, grouped as broader disease categories

We assessed the temporal ordering of the associations between Lyme disease (LD) and other diseases; specifically, we sought to determine whether a given comorbidity tended to occur before or after diagnosis of LD. We restricted our analysis to diseases with reported dates and only included the first reported encounter of a diagnosis, resulting in 41,713 disease-disease pairs, and analyzed their temporal ordering at the patient level. Specifically, for each comorbid disease pair (i.e. LD and another disease category), we tabulated the number of patients with both diseases and assessed which disease in the pair occurred first, or if they occurred at the same time, based on the visit dates. Overall, out of the 275 Lyme-comorbidity combinations for all time windows, 21 were nominally significant, with 5 diseases occurring prior to LD and 16 occurring after (*p* < 0.1; Table 1a). For the 2-year window, we identified three disease categories significantly associated with LD, with two prior and one after; for the 5-year window, 16 categories, with four prior and 12 after; and for the 10-year window, 17 categories with five prior and 12 after (Table 1a). We reconfirmed some previously reported Lyme comorbidities, including ‘nutritional deficiencies’ [p=0.069, probability (prob) =0.54 at 5 years; p=0.035, prob=0.55 at 10 years] ^30,31^, ‘vision defect’ (p=0.099, prob=0.57 at 5 years; p=0.076, prob=0.57 at 10 years) ^32 33^, and ‘disorder of lipid metabolism’ (p=0.036, prob=0.54 at 5 years; p=0.076, prob=0.57 at 10 years) ^34^. Additionally, we identified several disease comorbidities not previously reported, including ‘cataract’ (p=0.0037, prob=0.65 at 5 years; p=0.033, prob=0.64 at 10 years), ‘acute bronchitis’ (p=0.094, prob=0.62 at 2 years; p=0.022, prob=0.64 at 5 years; p=0.034, prob=0.63 at 10 years), and ‘nonmalignant breast conditions’ (p=0.057, prob=0.58 at 5 years; p=0.066, prob=0.58 at 10 years). A complete list of disease categories is shown in Table 1a.

**Table 1.**
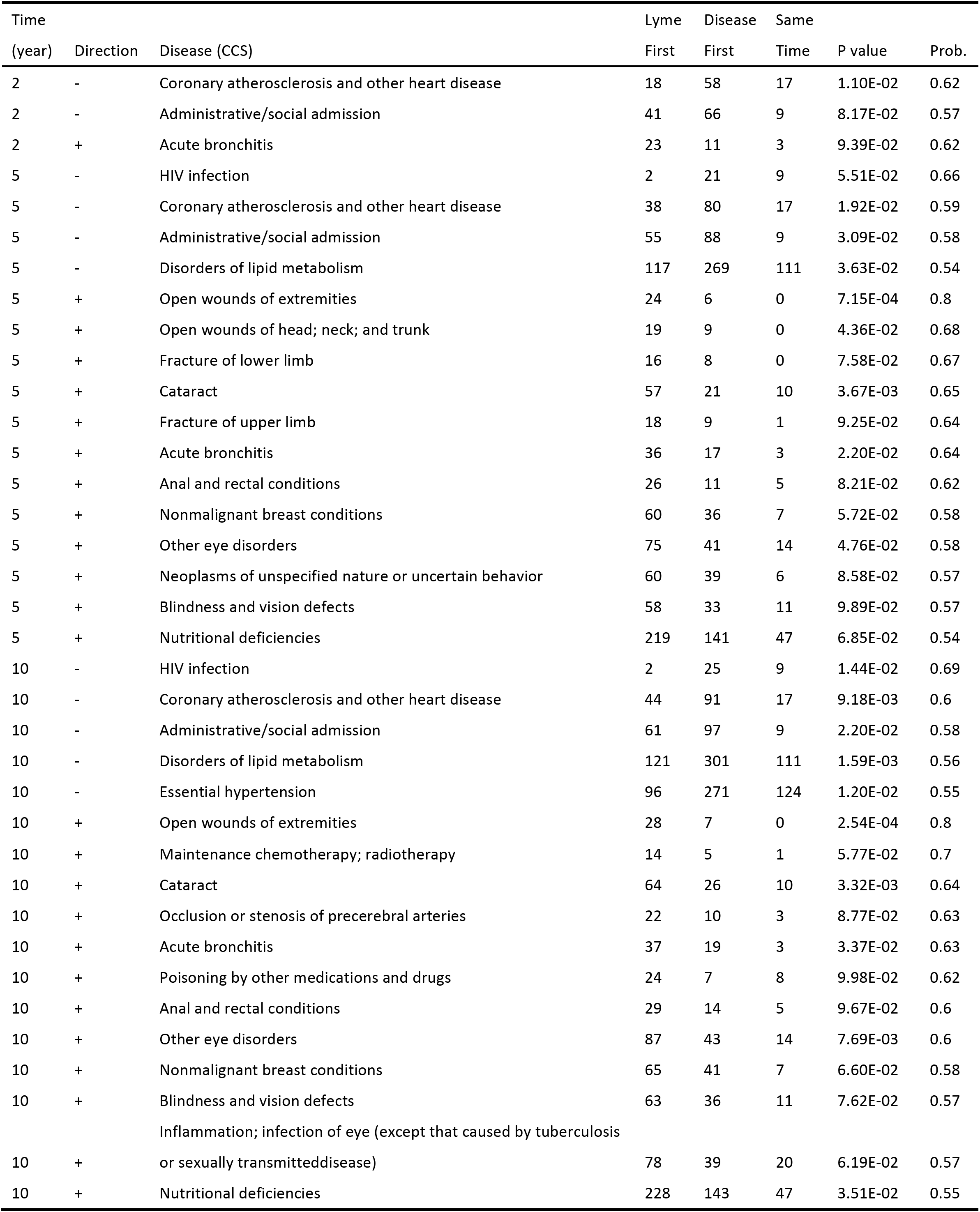
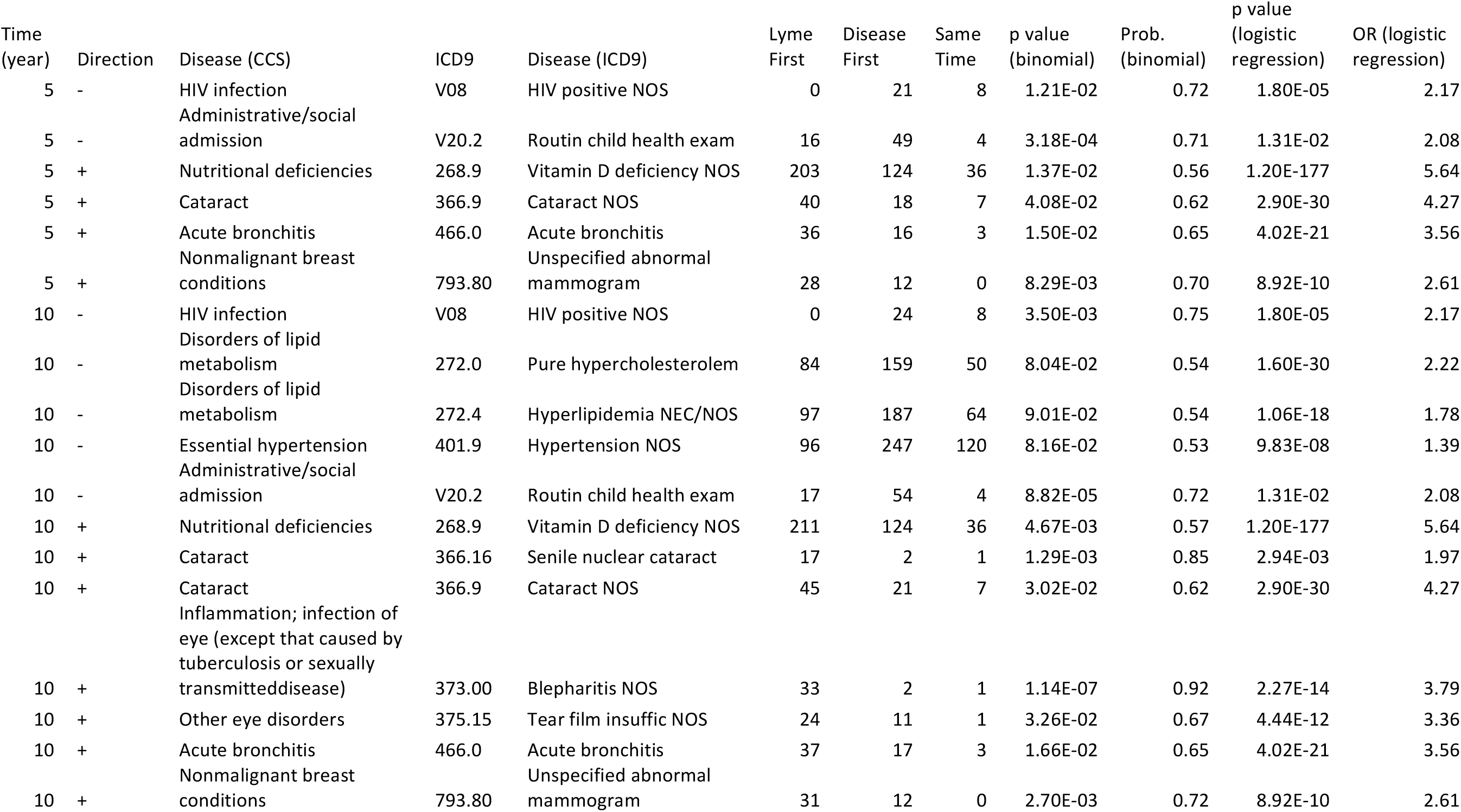
(a; top) Diseases associated with Lyme, analyzed as CCS-single-level categories (p value < 0.1). A total of 275 diseases were tested, (b; bottom) Diseases associated with Lyme, by ICD-9 category. A total of 3,639 diseases were tested. This table includes only the ICD-9 diseases that are classified into the CCS-single-level categories shown in (a), and a full list of associations is provided in Supplementary Table 1.

### Highlighted specific known or novel diseases associated with LD, analyzed at higher resolution

Although the CCS-single-level categories were helpful in identifying disease groups of relevance from a broader perspective, we also performed the same analysis at a higher granularity (Table 1b and S. Table 1). To this end, using the ICD-9 codes, we sought to determine which specific diseases drove the signal and whether the signal still persisted. A total of 3,639 Lyme–comorbidity combinations were analyzed using the ICD-9 codes. At the 2-year window, five pairs were nominally significant (p<0.1 due to the relatively small sample size, N≥20 patients for Lyme first onset), with four prior to LD and one after (S. Table 1). For the 5-year window, we found 53 significant associations, with 49 prior to LD and 4 afterwards. For the 10-year window, we found 75 significant associations, with 67 prior to LD diagnosis and 8 after. The significance of all disease categories significantly associated with LD that we identified in the previous analysis persisted, including the four diseases that significantly occurred prior to LD: ‘pure hypercholesterolemia’ (p=0.080, prob=0.54 at 10 years), ‘hyperlipidemia NEC/NOS’ (p=0.090, prob=0.54 at 10 years), ‘hypertension NOS’ (p=0.082, prob=0.53 at 10 years), and ‘coronary atherosclerosis (athero) NOS’ (p=0.022, prob=0.61 at 5 years; p=0.075, prob=0.62 at 10 years). Nine sequelae diseases, namely ‘vitamin D deficiency NOS’ (p=0.014, prob=0.56 at 5 years; p=0.0047, prob=0.57 at 10 years), ‘cataract NOS’ (p=0.041, prob=0.62 at 5 years; p=0.030, prob=0.62 at 10 years), ‘senile nuclear cataract’ (p=0.0013, prob=0.85 at 10 years), ‘tear film insufficiency (insuffic) NOS’ (p=0.033, prob=0.67 at 10 years), ‘acute bronchitis’ (p=0.015, prob=0.65 at 5 years; p=0.017, prob=0.65 at 10 years), ‘blepharitis NOS’ (p=1.1E-7, prob=0.92 at 10 years), ‘unspecified abnormal mammogram’ (p=0.0083, prob=0.70 at 5 years; p=0.0027, prob=0.72 at 10 years), ‘HIV positive NOS’ (p=0.012, prob=0.72 at 5 years; p=0.0035, prob=0.75 at 10 years)’ and ‘routine child health exam’ (p=3.2E-4, prob=0.71 at 5 years; p=8.8E-5, prob=0.72 at 10 years) drove the signal from the broad disease categories (Table 1b). Additionally, we reported LD comorbidities if other significant ICD-9 codes associated with LD with a list of comorbidities are not widely known, including ‘insomnia Not Elsewhere Classifiable (NEC)’ (p=0.042, prob=0.59 at 5 years; p=0.045, prob=0.58 at 10 years), ‘obstructive sleep apnea’ (p=0.085, prob=0.60 at 5 years; p=0.044, prob=0.62 at 10 years), ‘cervicalgia’ (p=0.073, prob=0.59 at 5 years; p=0.041, prob=0.60 at 10 years), and ‘dysuria’ (p=0.096, prob=0.59 at 5 years; p=0.034, prob=0.62 at 10 years).

We confirmed that the large majority of the comorbidity pairs were significantly associated with LD with concordant directionality by adjusting age, gender, and race by logistic regression (p<0.1). We provide a complete list of ICD-9 level disease associations that passed our significance threshold in both analyses Supplemental Table 1.

### Medications predicted to modulate risk of subsequent comorbidities in LD patients, analyzed as broader disease categories

To investigate how various prescribed medications influenced the risk of subsequent disease pathogenesis, we focused on comorbidities with onset after the first diagnosis of LD. Using the adaptive LASSO methodology and a logistic regression model, we investigated all medications prescribed to LD patients prior to the comorbidities. We found 3, 12, and 18 medications associated with disease comorbidities, classified by CCS-single-level categories, within 2, 5, and 10 years after Lyme diagnosis, respectively (S. Table 2, Figure 2a, Figure 2b). Four medication–Lyme comorbidity associations were supported by published studies ^35–41^, and we confirmed that these medications modulated the risks of Lyme comorbidities, including fluticasone–‘cataract’ (adjusted OR=1.94, p=0.072 at 5 years; adjusted OR=2.01, p=0.033 at 10 years) hydrochlorothiazide–’neoplasms of unspecified nature or uncertain behavior’ (adjusted OR=2.23, p=0.031 at 5 years; adjusted OR=2.48, p=0.0092 at 10 years), metformin–’nutritional deficiencies’ (adjusted OR=2.05, p=0.097 at 10 years), and esomeprazole–’nutritional deficiencies’ (adjusted OR=1.75, p=0.093 at 10 years).

**Figure 2.**
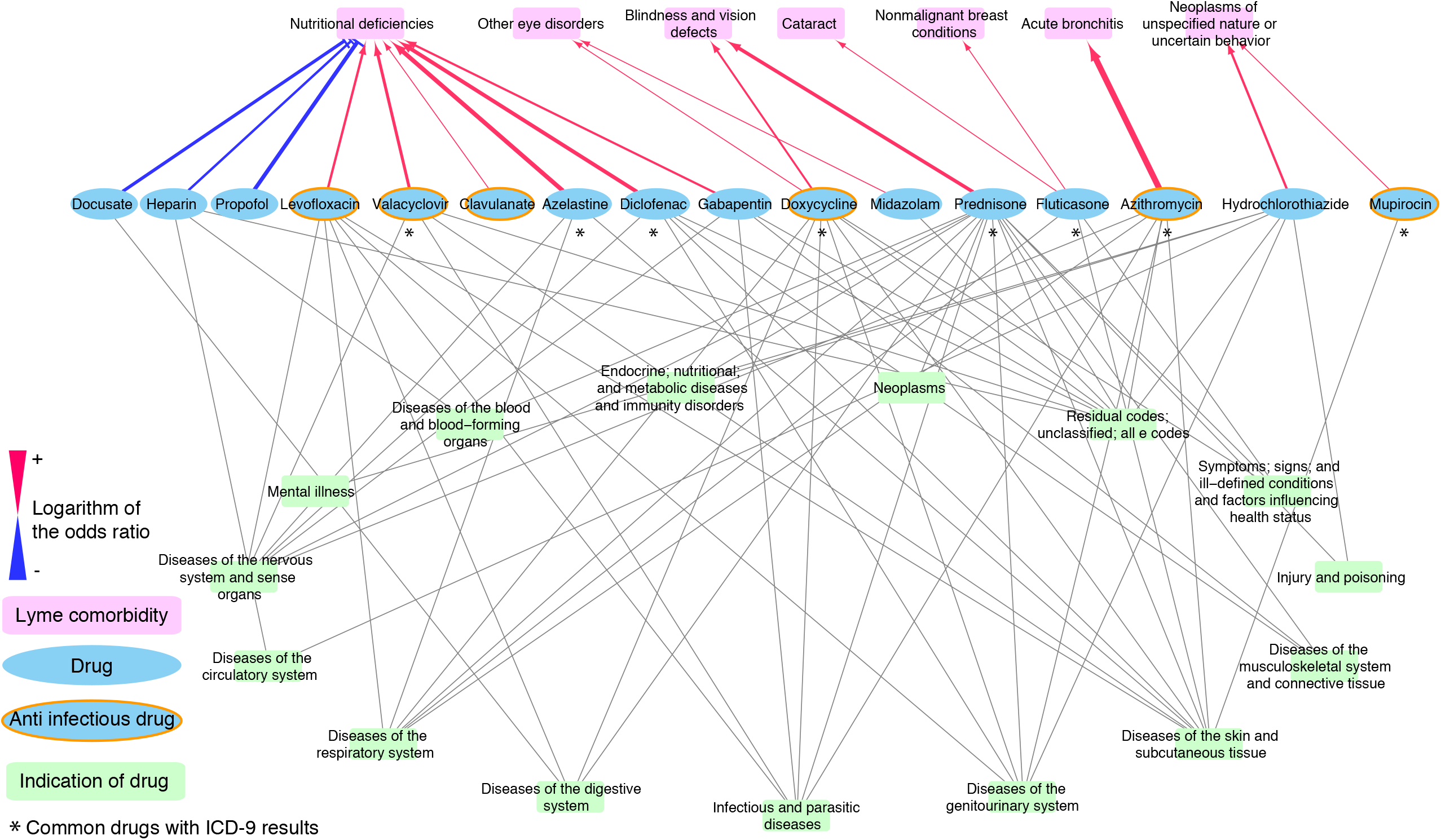
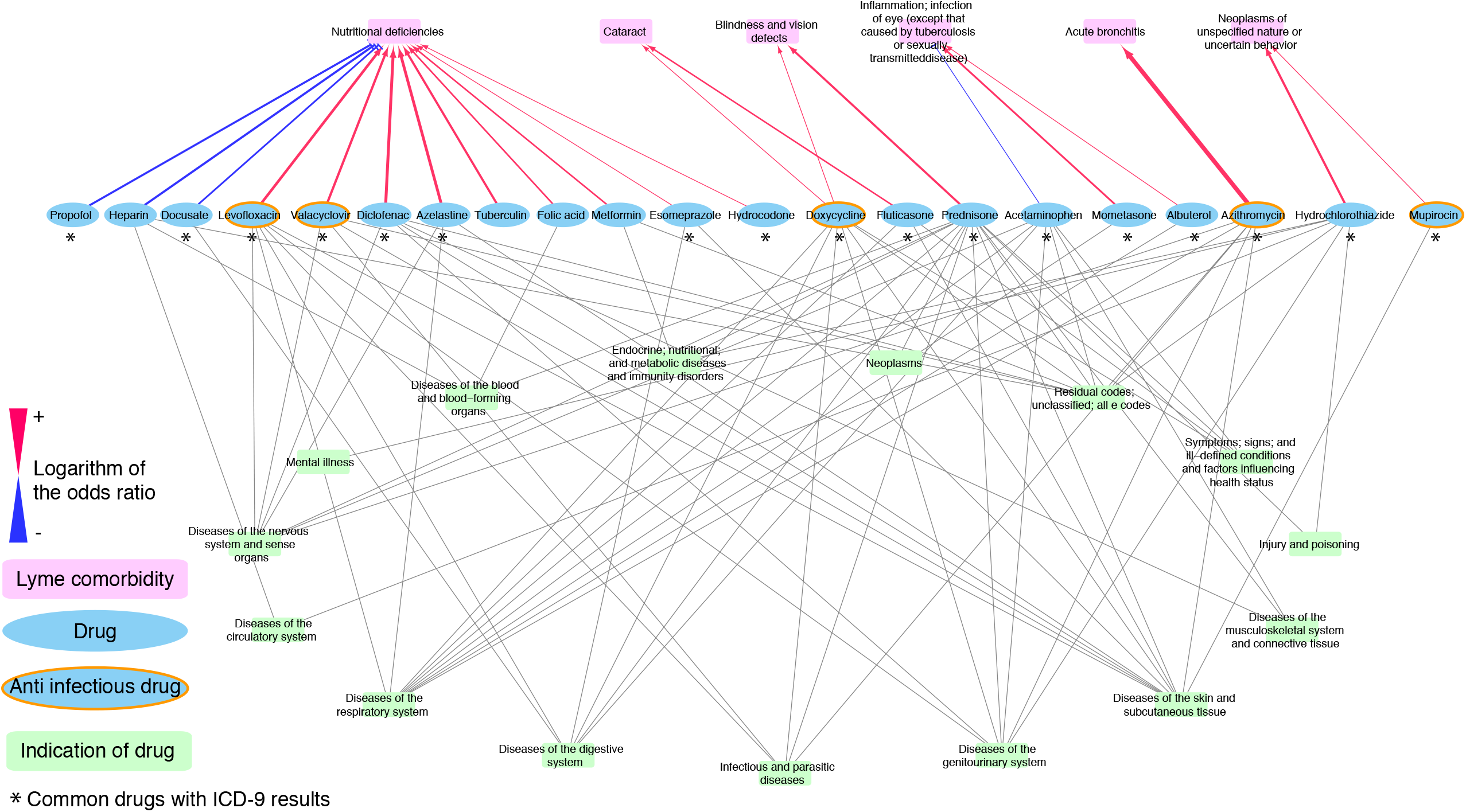
Medication–Lyme disease comorbidity network, analyzed by CCS-single-level categories, in time windows of 5 years (a) and 10 years (b). Significant associations between medications (cyan) and comorbidities (magenta) are connected by red or blue lines (p < 0.1). Red lines indicate risk associations (OR > 1), and blue lines indicate protective associations (OR < 1). Medications and indications (green) were connected based on information in the public knowledgebase MEDI ^60^.

Five antibiotics, doxycycline, azithromycin, levofloxacin, clavulanate, and mupirocin, and one antiviral drug, valacyclovir, were predicted to modulate the risk of subsequent comorbidities. Doxycycline, a first-line antibiotic that was the most prescribed antibiotic in our EMR for patients with LD (39%, N=553), was associated with an elevated risk of eye disorders, including ‘cataract’ (adjusted OR=2.05, p=0.092 at 2 years; adjusted OR=1.70, p=0.067 at 10 years), ‘blindness and vision disorders’ (adjusted OR=2.05, p=0.016 at the 5 years; adjusted OR=1.95, p=0.019 at 10 years), and ‘other eye disorders’ (adjusted OR=1.81, p=0.024) (Figure 2a).

In regard to ‘nutritional deficiencies’, eleven medications were predicted to be risk factors and three to be protective. Among the 11 risk factor medications were two antibiotics, levofloxacin (adjusted OR=2.26, p=0.0093 at 5 years; adjust OR=2.77, p=7.0E-4 at 10 years) and clavulanate (adjusted OR=1.64, p=0.094 at 10 years), and one antiviral prophylactic, valacyclovir (adjusted OR= 2.56, p=0.014 at 5 years; adjusted OR=2.58, p=0.011 at 10 years). Additionally, two pain relievers, diclofenac and hydrocodone, and one anti-allergy medication, azelastine, were also risk factors for ‘nutritional deficiencies’ (respectively: OR=3.08, p=0.0015 at 5 years/adjusted OR=3.02, p=0.0014 at 10 years; adjusted OR=1.90, p=0.034 at 10 years; and adjusted OR=3.44, p=0.0037/adjusted OR=2.95, p=0.0094). Interestingly, we could identify new therapeutic options for the LD adjunctive treatment. Three medications, propofol, docusate, and heparin, consistently exhibited a protective effect against ‘nutritional deficiencies’ at 5 and 10 years after LD (respectively: adjusted OR=0.32, p=6.6E-4 at 5 years/adjusted OR=0.43, p=0.0044 at 10 years; adjusted OR=0.37, p=0.0039 at 5 years/adjusted OR=0.46, p=0.014 at 10 years; and adjusted OR=0.44, p=0.024 at 5 years/adjusted OR=0.41, p=0.014 at 10 years) (Figure 2). In addition, acetaminophen exhibited a protective effect at the early stage (2 years post-Lyme) with adjusted OR=0.44, p=0.0069 (S. Table 2).

### Medications predicted to modulate risk of subsequent comorbidities in LD patients, analyzed at the ICD-9 level

In the higher-resolution analysis using ICD-9 codes, we identified 7, 22, and 31 medications that were significantly associated with the disease comorbidities at 2, 5, and 10 years post-Lyme (S. Table 3, Figure 3a and 3b). Among these were previously reported risk associations ^42–45^: for instance, steroid prednisone was a risk for ‘pain in limb’ with adjusted OR=2.16, p=0.030 at 5 years/adjusted OR=2.49, p=0.0063 at 10 years, and ciprofloxacin was a risk for ‘joint pain-shoulder (shlder)’ with adjusted OR=4.42, p=8.9E-5 at 5 years/ adjusted OR=4.39, p=3.1E-5 at 10 years. In addition, five of the side effects for four medications were reported in the SIDER database ^46,47^. Two steroids, fluticasone and mometasone, and one pain reliever, hydrocodone, were associated with increased risk for ‘acute upper respiratory infection (URI) NOS’ in comparison with the placebo group (S table 3) and were rediscovered in our study (respectively: adjusted OR=2.92, p=5.2E-5 at 5 years/adjusted OR=3.44, p=4.7E-7 at 10 years; adjusted OR=2.86, p=0.0028 at 10 years; adjusted OR=4.01, p=4.1E-5 at 5 years/adjusted OR=4.45, p=1.8E-6 at 10 years). We also reconfirmed the risk associations between fluticasone and ‘chronic rhinitis’ (adjusted OR=4.70, p=1.4E-4 at 2 years/adjusted OR=4.78, p=6.5E-7 at 5 years/adjusted OR=4.86, p=4.4E-8 at 10 years) and diclofenac and ‘pain in limb’ (adjusted OR=3.43, p=0.0011 at 10 years).

**Figure 3.**
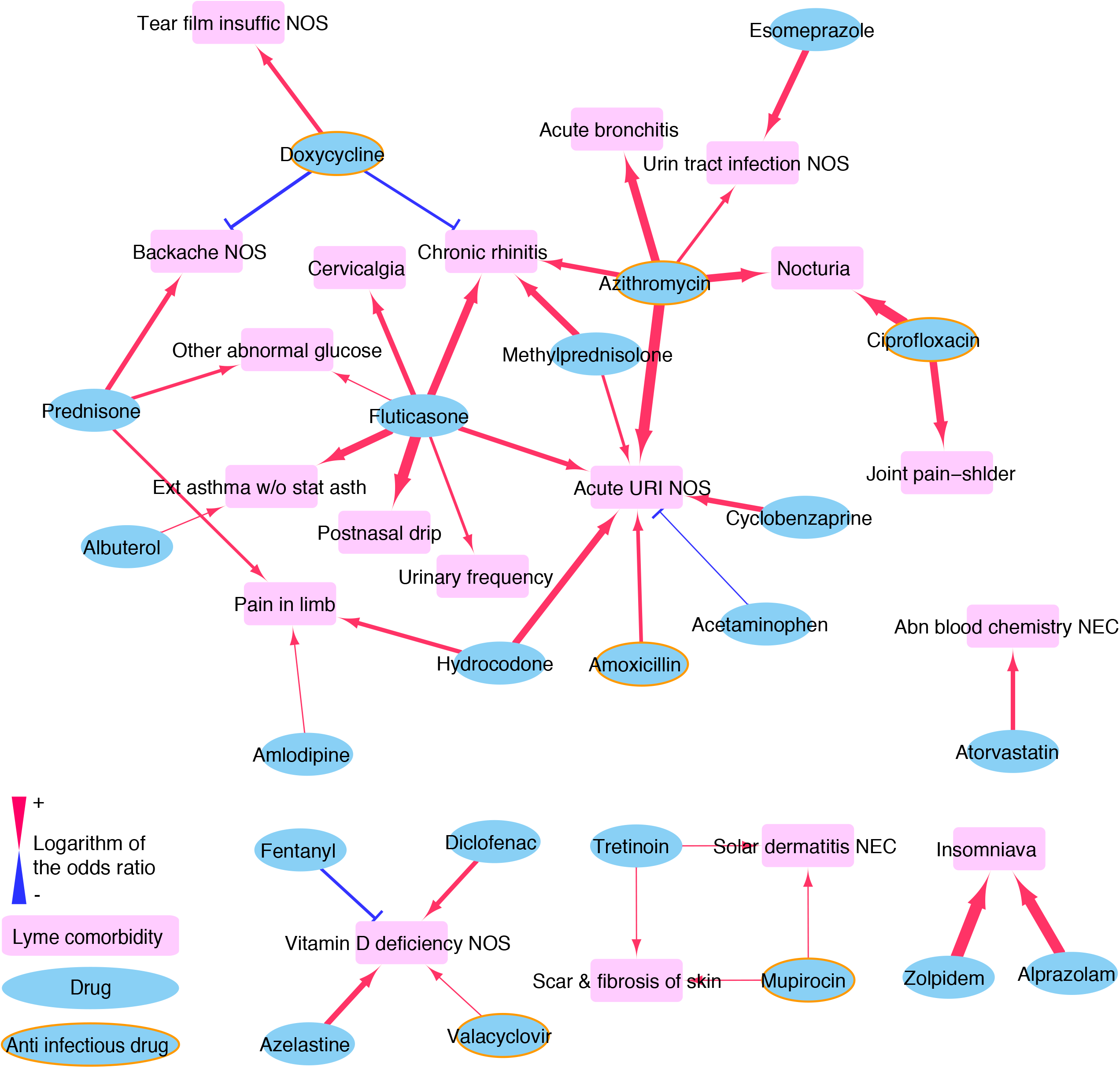
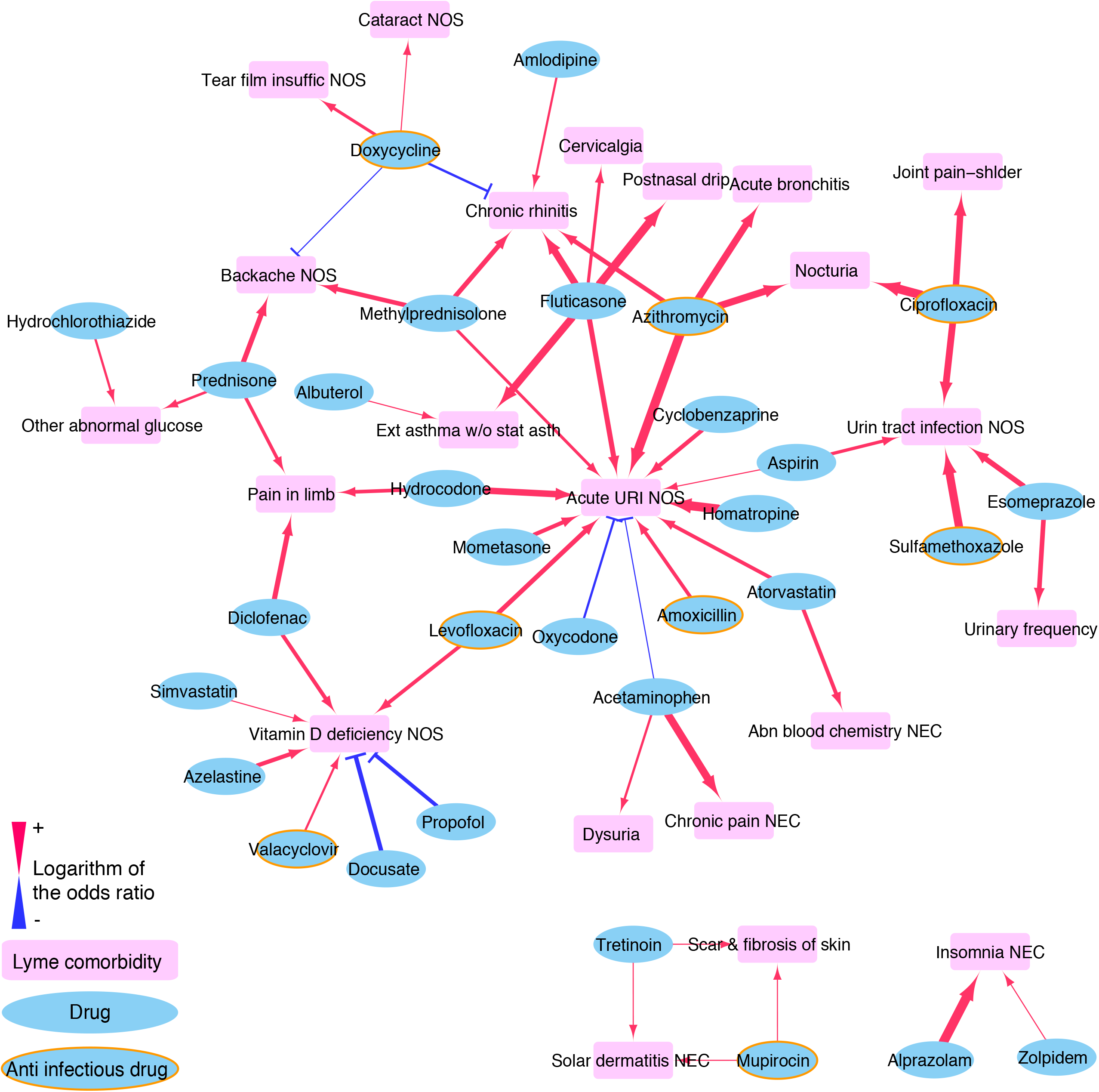
Medication–Lyme disease comorbidity network at the ICD-9 levels in time windows of 5 years (a) and 10 years (b). Significant associations between medications (cyan) and comorbidities (magenta) are connected by red or blue lines (p < 0.1). Red lines indicate risk associations (OR > 1), and blue lines indicate protective associations (OR < 1).

Doxycycline exhibited a consistently protective effect against typical symptoms of LD, including ‘backache NOS’ (adjusted OR=0.44, p=0.018 at 5 years/adjusted OR=0.50, p=0.035 at 10 years) and ‘chronic rhinitis’ (adjusted OR=0.48, p=0.036 at 5 years/adjusted OR=0.48, p=0.024 at 10 years) (Figure 3a, 3b). Furthermore, seven antibiotics, doxycycline, amoxicillin, azithromycin, ciprofloxacin, levofloxacin, mupirocin, and sulfamethoxazole, and one antiviral drug, valacyclovir, modulated the risk of subsequent comorbidities. Doxycycline consistently increased the risk of ‘cataract NOS’ (adjusted OR=2.57, p=0.053 at 2 years/adjusted OR=1.89, p=0.058 at 10 years), ‘tear film insuffic NOS’ (adjusted OR=2.64, p=0.042 at 5 years/adjusted OR=2.37, p=0.050 at 10 years), and ‘nocturia’ (adjusted OR= 3.46, p=0.010 at 2 years) (Figure 3b). Amoxicillin, another antibiotic recommended for LD, increased the risk of ‘acute URI NOS’ (adjusted OR=3.01, p=6.5E-4 at 2 years/adjusted OR=2.41, p=8.4E-4 at 5 years/adjusted OR=2.60, p=1.3E-4 at 10 years). Furthermore, azithromycin was associated with an increased risk of urinary-related diseases such as ‘nocturia’ (adjusted OR=4.62, p=3.5E-5 at 5 years/adjusted OR=4.90, p=1.3E-5 at 10 years) and ‘urinary (urin) tract infection NOS’ (adjusted OR=2.18, p=0.0095 at 5 years), as well as respiratory diseases such as ‘acute URI NOS’ (adjusted OR=8.63, p=3.9E-12 at 2 years/adjusted OR=7.55, p=1.0E-16 at 5 years/adjusted OR=7.2, p=1.4E-17 at 10 years), ‘acute bronchitis’ (adjusted OR=5.15, p=5.9E-5 at 5 years/adjusted OR=4.75, p=1.1E-4 at 10 years), and ‘chronic rhinitis’ (adjusted OR=2.99, p=7.2E-4 at 5 years; adjusted OR=3.07, p=1.5E-4 at 10 years). Ciprofloxacin, a fluoroquinolone antibiotic, increased risk of ‘nocturia’ (adjusted OR=7.19, p=5.9E-6 at 5 years/adjusted OR=6.92, p=7.0E-6 at 10 years) and ‘joint pain-shlder’ (adjusted OR=4.42, p=8.9E-5 at 5 years/adjusted OR=6.92 and p=7.0E-6 at 10 years). Mupirocin, an antibiotic used to treat skin infection, increased risk of skin disorders, including ‘solar dermatitis NEC’ (adjusted OR=12.93, p=6.9E-10 at 5 years/adjusted OR=13.96, p=3.0E-11 at 10 years) and ‘scar & fibrosis of skin’ (adjusted OR=11.52, p=6.9E-9 at 5 years/adjusted OR=12.46, p=1.0E-9 at 10 years) (Figure 3a, 3b).

‘Vitamin D deficiency NOS’, common in patients with persistent LD^48^, is a specific form of nutritional deficiency, a comorbidity identified earlier at the broader (CCS-single) level (Figure 2a, 2b). Several medications increased the risk of this condition, three at 5 years post-Lyme and five at 10 years. These included two anti-infective drugs, levofloxacin (adjusted OR=2.68, p=0.0012 at 10 years) and valacyclovir (adjusted OR=1.95, p=0.087 at 5 years/adjusted OR=2.01, p=0.065 at 10 years), azelastine (adjusted OR=3.37, p=0.0035 at 5 years/adjusted OR=2.92, p=0.0085 at 10 years), diclofenac (adjusted OR=2.75, p=0.0032 at 5 years/adjusted OR=2.75, p=0.0026 at 10 years), and simvastatin (adjusted OR=1.77, p=0.083 at 10 years). On the other hand, four medications protected against ‘vitamin D deficiency NOS’: docusate (adjusted OR=0.33, p=0.0015 at 10 years), propofol (adjusted OR=0.37, p=0.0015 at 10 years), fentanyl (adjusted OR=0.46, p=0.012 at 5 years), and acetaminophen (adjusted OR=0.39, p=0.0025 at 2 years) (Figure 3a, 3b).

Respiratory disease (Figure 3b) is a complication frequently reported after LD ^49^. We identified 11 medications that increased risk for these conditions and two that exhibited protective effects. In addition to the three medications reported in SIDER database and amoxicillin and azithromycin above, the medications that conferred increased risk for ‘acute URI NOS’ were an antibiotic, levofloxacin (adjusted OR=3.18, p=9.1E-4 at 10 years), a steroid, methylprednisolone (adjusted OR=2.14, p=0.027 at 5 years/adjusted OR=2.31, p=0.0077 at 10 years), cyclobenzaprine (adjusted OR=3.21, p=0.0025 at 5 years/adjusted OR=2.91, p=0.0037 at 10 years), homatropine (adjusted OR=7.16, p=7.3E-7 at 10 years), atorvastatin (adjusted OR=2.67, p=0.0017 at 10 years), and aspirin (adjusted OR=1.80, p=0.073 at 10 years). By contrast, acetaminophen (adjusted OR=0.55, p=0.025 at 5 years/adjusted OR=0.62, p=0.050 at 10 years), and oxycodone (adjusted OR=0.49/p=0.021 at 10 years) were associated with protective effects against this disease.

### Medications that modulate LD pathophysiology on different timescales

We identified 16 medications associated with disease comorbidities within 5 years post-LD, 81% (13/16) of which overlapped with those identified as associated 10 years post-Lyme by the CCS-single-level categorization. Specifically, five out of six anti-infective drugs, doxycycline, azithromycin, levofloxacin, mupirocin, and valacyclovir, appeared in both timeframes.

Moreover, 22 medications were associated with the ICD-9–level disease comorbidities within 5 years after Lyme, 95% (21/22) of which were also identified in the 10-year post-Lyme analysis. Among those 21 medications, five are antibiotics (doxycycline, amoxicillin, azithromycin, ciprofloxacin, mupirocin) and one is an antiviral drug (valacyclovir). The four medications associated with comorbidities exclusively in the 5 years post-Lyme, clavulanate, gabapentin, midazolam, and fentanyl, may impact relatively early Lyme comorbidities (S. Figure 1).

A total of 17 medications overlapped between the CCS-single and ICD-9 levels in either the 5-year or 10-year time windows. Five of them were anti-infective drugs, namely doxycycline, azithromycin, levofloxacin, mupirocin, and valacyclovir. In the 5-year time window, 16 medications were associated with comorbidities classified by CCS-single-level category, of which 50% (8/16) were also identified at the ICD-9 level. At 10 years post-Lyme diagnosis, we identified 21 significant associations between medications and comorbidities, of which 81% (17/21) were consistent with those identified at the ICD-9 level.

### Survival analysis of first-line medications in propensity-matched populations

By the cross-sectional analysis described above, we demonstrated that certain medications increased risk or protected against disease comorbidities in patients with LD. At higher resolution (i.e., using ICD-9 codes) with 10-year follow up, we found that doxycycline, the most commonly used antibiotic for treatment of LD ^13^, protected against ‘backache NOS’ and ‘chronic rhinitis’, but increased risk of ‘tear film insuffic NOS’ and ‘cataract NOS’. Another commonly used antibiotic, amoxicillin, was associated with elevated risk of ‘acute URI NOS’.

Prior to propensity score matching, we identified significant differences in the distributions of demographic and clinical characteristics between the doxycycline/amoxicillin-treated and untreated groups before. The doxycycline-treated group was significantly older than the untreated group (P < 0.007), whereas the amoxicillin-treated group was significantly younger than the untreated group (P=8.7E-4). In addition, doxycycline was prescribed more frequently to male than female patients (P < 0.03). The doxycycline/amoxicillin-treated groups had higher prevalences of certain pre-existing comorbidities and a higher prescription rate of particular medications than the untreated groups (S. Table 4). Moreover, both the doxycycline/amoxicillin treated groups had higher propensity scores than the corresponding untreated groups (P < 0.001). To clarify the longitudinal effects of doxycycline and amoxicillin, we analyzed these associations by propensity-score-matched survival analyses. After propensity score matching, the control cohorts were well balanced with the treated groups in terms of observed covariates (S. Table 4).

This analysis revealed that the risk of ‘backache NOS’ (Figure 4a) and ‘chronic rhinitis’ (Figure 4b) was significantly lower in the doxycycline-treated cohort than in the untreated cohort (HR=0.42, p=0.020; HR=0.49, p=0.040, respectively; Table 3). Furthermore, Kaplan-Meier curves demonstrated that the cumulative probabilities of remaining free from ‘cataracts NOS’ and ‘tear film insuffic NOS’ were lower among doxycycline-treated patients (p = 0.0672 and 0.0608, respectively; Figures 4c and 4d). Cox regression analysis supported a statistically significant association between doxycycline usage and increased risk of both ‘cataract NOS’ and ‘tear film insuffic NOS’ (HR=1.90, p=0.072; HR=2.65, p=0.071). On the other hand, patients prescribed amoxicillin had significantly higher hazard ratios for ‘acute URI NOS’ (HR=2.26, p=0.0091; Figure 4e). Therefore, the effects of doxycycline and amoxicillin revealed by the cross-sectional analysis were confirmed by survival analyses using the propensity score-matched cohort (Table 3).

**Figure 4.**
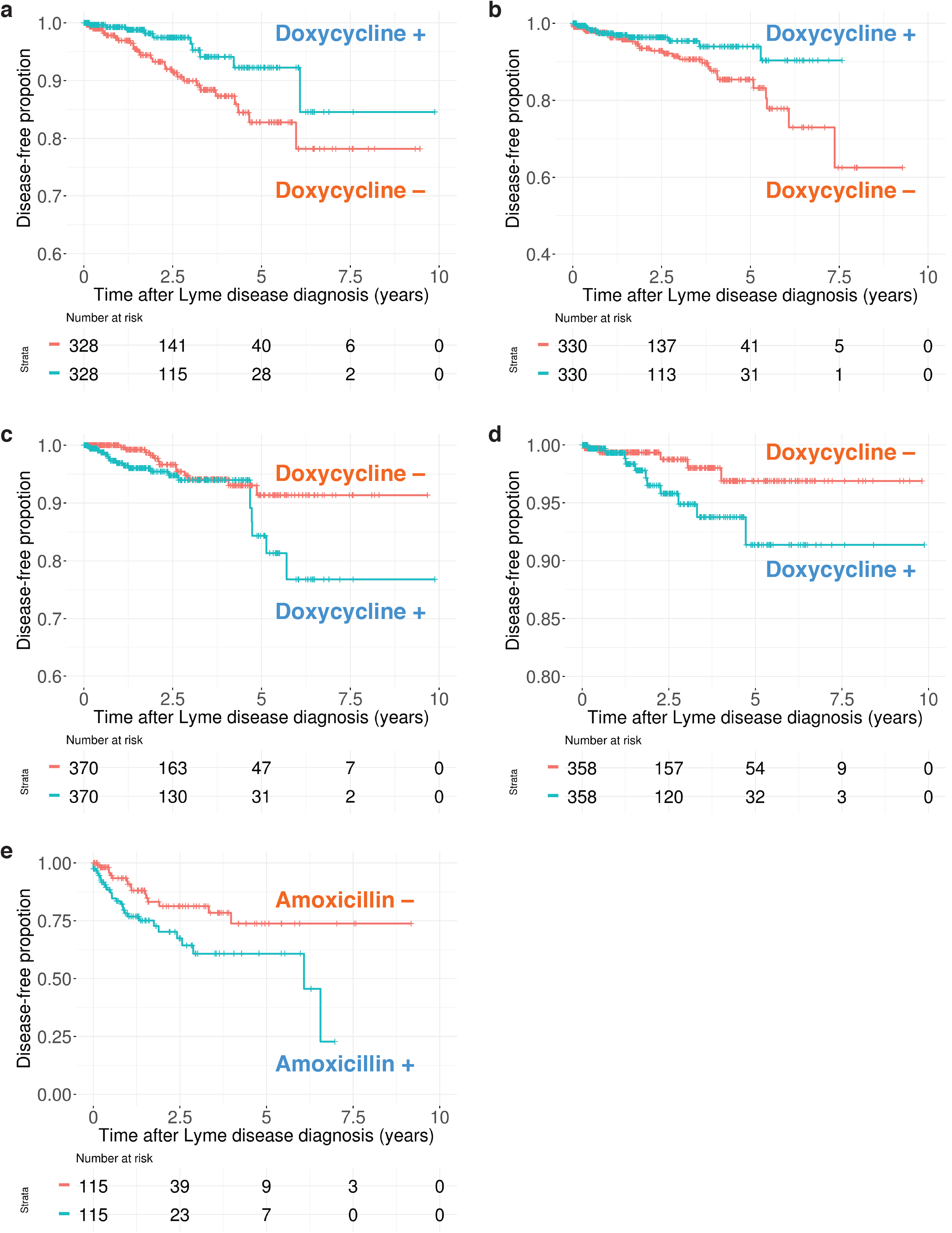
Kaplan–Meier plot of propensity-score-matched survival analysis (a) doxycycline–‘backache NOS’ (ICD-9 code: 724.5), (b) doxycycline–’chronic rhinitis’ (472.0), (c) doxycycline–‘cataract NOS’ (366.9), (d) doxycycline-’tear film insuffic NOS’ (375.15), and (e) amoxicillin–‘acute URI NOS’ (465.9).

**Table 2.**
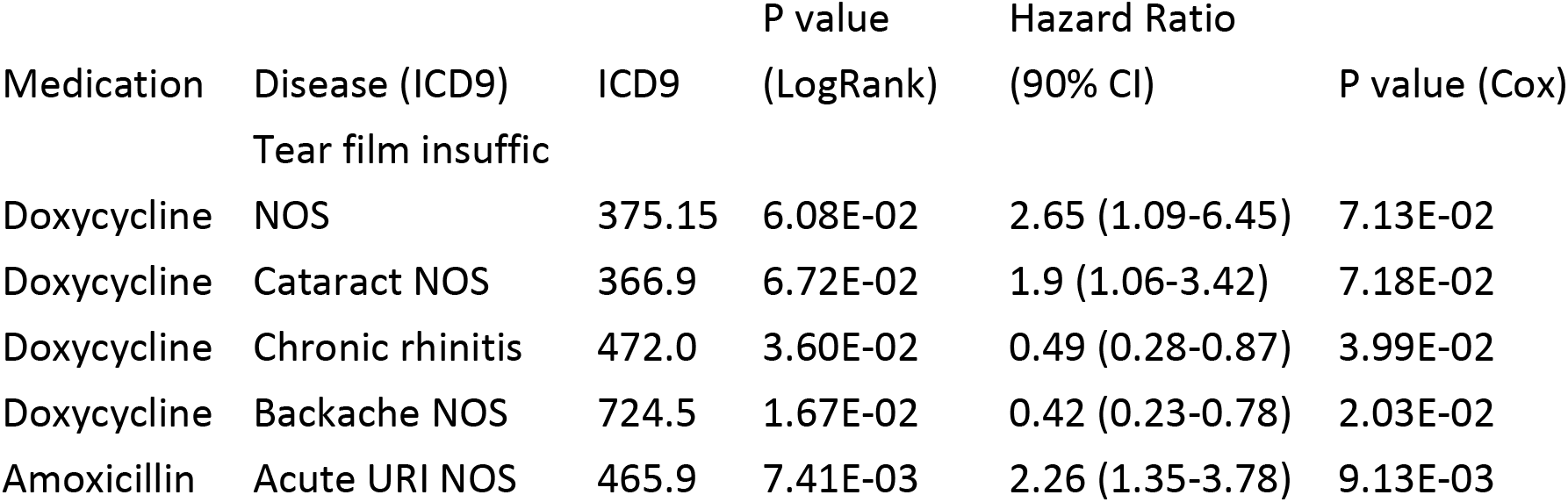
Survival analyses of first-line therapeutics for Lyme disease using a propensity-score-matched cohort.

## DISCUSSION

Proper diagnosis, treatment, and management of Lyme Disease (LD) are difficult for a variety of reasons. In particular, the complex interplay between various treatments and current clinical status, including disease burden, can lead to a wide range of sequelae. This study represents the first data-driven effort to identify clinical factors that affect treatment of LD patients using large-scale EMR data. EMR systems contain information pertaining to patients’ health over time, including treatments administered and clinical outcomes. Thus, one advantage of using these data in comparison with prospective clinical trial studies is the availability of longitudinal data spanning more than 10 years, increasing the likelihood of capturing long-term effects. We utilized statistical and machine learning analyses to identify associations between medications, including first-line treatments, and comorbidity pathogeneses. In contrast to one-size-fits-all strategies, our approach may facilitate the personalization of treatment regimens based on the clinical profiles of affected individuals. This strategic transition is essential in light of the tremendous variability in efficacy of antibiotics and the adverse events associated with these treatments.

We first identified all significant comorbidities of patients with LD before and after their Lyme infections. Next, we applied machine learning models to assess the effect of medication treatment on the risk of developing subsequent conditions. Our analyses identified known associations between medications and specific disease comorbidity outcomes, and also discovered connections between drugs and LD that could facilitate precision medicine aimed at tailoring treatments to affected individuals’ Lyme symptom profile.

Our analysis identified co-morbid conditions that were typically present before LD infection. Although this disease clearly requires contact with the bacteria, certain physiological properties make individuals more or less susceptible to infection. For example, we found that individuals categorized as having ‘disorders of lipid metabolism’ were more likely have LD infection in the future. *Borrelia burgdorferi* requires cholesterol for growth; researchers have found that apolipoprotein E (apoE)-deficient and low-density lipoprotein receptor (LDLR)-deficient mice, which have high levels amounts of serum cholesterol, are more susceptible than wild-type mice to pathogenesis induced by this bacterium ^34^. Additionally, patients with hypercholesterolemia could increase susceptibility to trigger tick bite for this vector borne disease due to body heat, CO2, and moisture which are key attractants similarly to mosquitoes ^50 51^, but this is beyond the scope of this analysis and needs to be further investigated. Additionally, we identified an association between HIV infection and LD. Specifically, our data showed that patients with HIV almost exclusively developed LD subsequent to the HIV diagnosis, suggesting that immune system alteration increases the risk of LD. A handful of case reports have indicated that HIV-positive immunocompromised patients develop more severe Lyme complications following infection ^52–54^. Although the specific immunological mechanisms driving this connection remain unclear, it seems reasonable to speculate that immunosuppression plays an important role ^55^.

We also identified conditions that are more likely to be present after a LD diagnosis than beforehand, consistent with the possibility that these diseases are side effects or complications arising from Lyme infection. Many of these associations are well documented, enhancing our confidence in our results. Specifically, we found that the categories ‘nutritional deficiencies’, ‘cataract’, ‘acute bronchitis’, ‘other eye disorders’, and ‘Inflammation; infection of eye (except that caused by tuberculosis or sexually transmitted disease)’ are more likely to occur after Lyme infection than beforehand or by chance.

The results of this analysis feed into our drug-comorbidity associations network and can be used to inform treatment regimens. The associations identified for first-line antibiotic treatments of LD have the most straightforward potential application. These agents act by killing *B. burgdorferi*, and thus prevent the development of many complications associated with prolonged exposure to the bacteria. However, while antibiotics are the most effective first-line treatments for LD, their efficacies are nonetheless limited; moreover, some symptoms may persist notwithstanding the use of antibiotics, and long-term exposure to these agents risks additional complications. Indeed, it is possible that even acute use (usually a month) of these treatments is associated with long-term complications of LD that yet to be determined. Even when treated, up to 20% of patients develop Post-Treatment Lyme Disease Syndrome (PTLDS), in which symptoms including fatigue or muscle pain last for months or years. Although the etiology of PTLDS is not yet known, better tailoring of treatment strategies to an individual’s phenotypic profile could prevent or modulate the risk of developing these symptoms. We identified a number of comorbidities matching the symptoms aligned with PTLDS, including chronic pain, chronic rhinitis (Figures 3a and 3b). Notably in this regard, we also found that usage of steroid medications increases the risk of many symptoms common to PTLDS. Consistent with this, corticosteroid use is associated with poor outcomes for LD patients ^43^. In particular, we found that prednisone was associated with elevated risk for ‘backache NOS’, ‘pain in limb’ and ‘other abnormal glucose’, defined at the ICD-9 level. Use of fluticasone was associated with elevated risk for ‘chronic rhinitis’, ‘postnasal drip’, ‘cervicalgia’, and ‘acute URI NOS’; mometasone was also associated with elevated risk for ‘acute URI NOS’; and methylprednisolone was linked to increased risk of ‘chronic rhinitis’, ‘backache NOS’, and ‘acute URI NOS’ as well. Steroids, which suppress patients’ immune systems, might be particularly harmful to LD patients, allowing the bacteria to grow, rather than attacking the infections. These findings suggest that steroid use should be limited in LD patients, and that patients exposed to these drugs should be monitored carefully for complications.

Other findings from our drug-comorbidity network might facilitate personalization of treatment regimens, with more favorable clinical outcomes for patients. For example, several anti-infectious drugs, pain relievers such as diclofenac and hydrocodone, and the anti-allergy medication azelastine were also associated with higher rates of ‘nutritional deficiencies’, suggesting that physicians should consider recommending vitamin supplements for patients receiving these treatments.

Doxycycline is already associated with a range of side effects, including pain, increased pressure inside the skull ^56^, and gastrointestinal injury ^57^. The nuances of these associations are not well understood. On the other hand, we also found that doxycycline use was associated with lower risk of ‘backache NOS’ and lower rates of ‘chronic rhinitis’; the latter is to be expected, as it is a common symptom of chronic LD ^11^. This medication increased the risk of many eye-related issues, such as ‘cataract NOS’, with the 2- and 10-year time windows, suggesting that it might exert both short-term and long-term side effects. We also found associations between this medication and elevated risk for ‘tear film insuffic NOS’ and ‘blindness and vision disorders’.

The development of respiratory-related complications is a major concern for patients infected with LD. In one case study, secondary adult respiratory distress syndrome caused the death of a patient affected with LD during the course of her 2-month treatment ^58^. The patient did not respond to conventional treatments, including antibiotics, salicylates, and steroids. In our study, we identified medications that are associated with elevated risk of respiratory-related diseases. Specifically, we found that the antibiotics amoxicillin, levofloxacin, and azithromycin all conferred increased risk of ‘acute URI NOS’. In addition to the known risks of two steroids, prednisolone, and mometasone, reported in the SIDER database (S table 3), we found that another steroid, methylprednisolone, was also associated with increased risk for this disease. While we are unable to infer causation from our analyses, the associations we identified will hopefully inform physicians of the risks and encourage them to take the appropriate prophylactic measures. Particular attention should be paid to Lyme patients with respiratory complications because their immune systems are already weakened from Lyme infection, and certain medications (such as amoxicillin) are ineffective at treating these infections ^59^.

This study had several limitations. One issue is the relative low sample size, which is a consequence of the rarity of this disease although our hospital has the largest EMR system in NYC. Based on the available ELISA and Western blot lab tests specific for IgM and IgG, we found a great concordance for positive serology in patients with Lyme ICD codes, which enhanced our confidence of identifying true positive LD patients. Based on limited availability of these data, however, we had to use the ICD-9 code alone to select the patient cohort in this study. As tick bites are most likely to occur in surrounding rural locations in which forests are present, many patients may be initially diagnosed in a different facility, and then come to MSH for follow-up treatment. Another limitation is related to the close proximity of MSH to other medical centers in the area. Specifically, patients may seek treatment at other nearby hospitals, resulting in the loss of valuable information from our EMR system. Finally, because we do not have access to patients’ historic EMR data from outside of MSH, our temporal analyses may not accurately capture the true timeline of acquisition of disease comorbidities. We are currently performing an external replication analysis at another academic medical center, and the results of this effort may bolster our conclusions. Additionally, we are applying the findings from our current study in order to model explicit, optimal treatment recommendations at the patient level. From this work, we hope to enhance not only the success rates of treatment of LD, but also to facilitate preemptive strategies for managing high-risk ensuing conditions.

Our study is the first to investigate a comprehensive and racially diverse EMR with the aim of discovering the detailed clinical profiles of patients before and after diagnosis of LD. We identified a list of medications, including antibiotics recommended for treatment, which represent possible risk factors for chronic LD or PTLDS. In addition, we hope to investigate the contributions of genomics and genetic variants to differences pathophysiology. Our predictive medication–comorbidity models provide an evidence-based approach for treatment regimens that takes into account risk for Lyme comorbidities, with the ultimate goal of guiding precision medicine based on the individual clinical phenotypes of patients with LD.

## CONTRIBUTORS

LL and JD were responsible for initial study design. LL was responsible for data collection. OI, BG, and LL were responsible for study implementation, interpreting the data, and ongoing management. OI and BG were responsible for statistical analysis, predictive models, literature review, and generating tables and figures. OI, BG, LL, and JD and wrote and edited manuscript. BK provided HIV insights and edits. LL and JD supervised the study.

## ACKNOWLEDGEMENTS

This study was funded by the Steven & Alexandra Cohen Foundation. We thank Savi Glowe and Francisco Santiago for administrative support. We thank Dr. Christopher Patil for providing scientific edits. We thank Mount Sinai Data Warehouse for supporting data access.

## Figure legends

**Supplementary figure 1** Venn diagram of the medications that significantly associated with at least one disease comorbidity in the 5- and 10-year time windows. (a) CCS-single-level categories. (b) ICD-9 level.

**Supplementary table S1** All diseases associated with Lyme, by ICD-9 category (p value < 0.1). **Supplementary table S3** Medications predicted to modulate risk of disease comorbidities, by CCS-single-level category (p value < 0.1).

**Supplementary table S4** Medications predicted to modulate risk of disease comorbidities, by ICD-9 category (p value < 0.1).

**Supplementary table S5** The balance of covariates before and after propensity score matching.

